# Ring-finger protein 34 facilitates nervous necrosis virus evading antiviral innate immunity by targeting TBK1 and IRF3 for ubiquitination and degradation

**DOI:** 10.1101/2022.12.05.519093

**Authors:** Wanwan Zhang, Leshi Chen, Lan Yao, Peng Jia, Yangxi Xiang, Meisheng Yi, Kuntong Jia

**Author notes:** Corresponding author (Meisheng Yi); (Kuntong Jia).

## Abstract

Ubiquitination, as one of the most prevalent posttranslational modifications of proteins, enables a tight control on host immune responses. Many viruses hijack the host ubiquitin system to regulate host antiviral responses for their survival. Here, we found that fish pathogen nervous necrosis virus (NNV) recruited an E3 ubiquitin ligase ring finger protein 34 (RNF34) to inhibit RLRs-mediated interferons (IFN) response via ubiquitinating TBK1 and IRF3. Ectopic expression of RNF34 greatly enhances NNV replication and prevents IFN production, while deficiency of RNF34 led to the opposite effect. Furthermore, RNF34 targets TBK1 and IRF3 via its RING domain. Of note, the interactions between RNF34 and TBK1 or IRF3 were conserved in different fish species. Mechanically, RNF34 promote K27-linked ubiquitination and degradation of TBK1 and IRF3, which in turn diminishing TBK1-induced translocation of IRF3 from cytoplasm to nucleus. Ultimately, NNV capsid protein (CP) was found directly bind with RNF34 and this interaction was conserved in different fishes, and CP induced TBK1 and IRF3 degradation and IFN suppression was depended on RNF34. Our finding demonstrated a novel mechanism by which NNV CP evaded host innate immunity via RNF34, and provided a potential drug target for the control of NNV infection.

**Author Summary:** Ubiquitination plays an essential role in the regulation of innate immune responses to pathogens. NNV, a kind of RNA virus, is the causal agent of a highly destructive disease in a variety of marine and freshwater fish. Previous study reported NNV could hijack the ubiquitin system to manipulate the host’s immune responses, however, how NNV utilizes ubiquitination to facilitate its own replication is not well understood. Here, we identified a novel distinct role of E3 ubiquitin ligase RNF34 as an IFN antagonist to promote NNV infection. Nervous necrosis virus capsid protein utilized RNF34 to target TBK1 and IRF3 for K27 and K48-linked ubiquitination degradation. Importantly, the interactions between RNF34 and CP, TBK1 or IRF3 are conserved in different fishes, suggesting it is a general immune evasion strategy exploited by NNV to target the IFN response via RNF34.

## Introduction

Ubiquitination is a protein modification occurring post-translationally that conjugating the 76-amino acid polypeptide ubiquitin to substrate proteins through lysine residues (K6, K11, K27, K29, K33, K48, and K63) [1]. A cascade of enzymes are responsible for ubiquitination, including ubiquitin-activating enzymes (E1), ubiquitin-conjugating enzymes (E2), and an ubiquitin ligases (E3), among which E3 ubiquitin ligases are of particular interest due to their substrate specificity [2]. Based on the presence of specific functional domains and the mechanism of catalysis, E3 ubiquitin ligases are divided into three major classes, including RING, RING-between-RING and HECT E3 ubiquitin ligase [3]. Numerous studies have demonstrated that E3 ubiquitin ligases play important roles in a variety of biological and cellular processes, including but not limited to protein trafficking, apoptotic cell death, innate immune responses and virus infection. For example, E3 ubiquitin ligase deltex-4 (DTX4) is recruited by NLRP4 and ubiquitinates TBK1 at K48-linked chains, thereby inhibiting interferon (IFN) signaling [4]. TRIM40 binds both MDA5 and RIG-I, and then promotes their polyubiquitination degradation through K27- and K48-linked chains, leading to a strong limitation on IFN production [5].

The innate immunity, as the first host defense line, would be initiated following the sense of viral infection. RIG-I-like receptors (RLRs), responsible for the recognition of RNA virus, evokes a downstream signaling cascade and then activates TANK-binding kinase 1 (TBK1) [6]. The activated TBK1 further motivates interferon regulatory factor 3 (IRF3) and promotes IRF3 translocation into the nucleus, which finally induces IFN and a series of interferon-stimulating genes (ISGs) production [7]. To maintain host immune homeostasis, strict and precise immune system regulation is essential. Several studies have demonstrated that E3 ubiquitin ligases act as key regulators of the RLR-signaling pathway. For example, TBK1 undergoes posttranslational modifications including K63 or K48-linked polyubiquitination mediated by TNF receptor-associated factor 3 (TRAF3) or RNF128 for IFN signaling optimization during virus infection [8, 9]. RNF153 promotes the K48-linked ubiquitination degradation of mitochondrial antiviral signaling protein (MAVS) aggregates, suppressing MAVS-mediated IFN signaling [10].

Accumulating evidence shows that RING-type E3 ubiquitin ligases, the largest class of E3s, have been associated with the regulation of many aspects of the immune system [11]. For instance, RNF122 binds to RIG-I to induce K48-linked ubiquitination degradation of RIG-I, leading to the inhibition of type I IFN production [12]. RNF166 binds to TRAF3 and tumor necrosis factor (TNF)-associated factor 6 (TRAF6) and promotes the ubiquitination of TRAF3 and TRAF6 to enhance IFN-β production [13]. As a response to antiviral state in infected cells, many E3 ubiquitin ligases were hijacked by viruses to counteract the immune response [14]. For instance, hepatitis B e antigen suppressed the TRAF6-dependent K63-linked ubiquitination of NEMO, thereby downregulating nuclear factor kappa B (NF-κB) activity and promoting virus replication [15].

Nervous necrosis virus (NNV), belonging to the member of the genus *Betanodavirus* in *Nodaviridae* family, is a fish RNA virus that is prevalent worldwide and results in up to 100% mortality in affected larvae and juvenile fish [16]. NNV infection has caused considerable economic losses in aquaculture. NNV consists of two molecules of positive-sense single-stranded RNA (RNA1 and RNA2), which encodes RNA dependent RNA polymerase (RNA1) and capsid protein (CP, RNA2), respectively. Recently, we and other scholars reported that NNV could hijack the ubiquitin system to manipulate the host’s immune responses. For example, NNV CP induced polyadenylate binding protein degradation to stimulate host translation shutoff by the ubiquitin-proteasome system [17]. We found that capsid protein of NNV targeted TRAF3 and IRF3 for ubiquitination and degradation, leading to the suppression of IFN production. Particularly, RNF114 was utilized by CP to promote ubiquitination and degradation of TRAF3 [18]. However, which E3 ubiquitin ligase is responsible for CP induced ubiquitination of IRF3 remains unknown.

In this study, we identified that RNF34 as the suppressor of RLRs-mediated type I IFNs production during NNV infection. RNF34 interacted with TBK1 and IRF3 and promoted their ubiquitination degradation. Furthermore, CP recruited RNF34 to evade host innate immunity. Our findings identified an evasion strategy employed by NNV to evade RLRs-mediated antiviral immune responses via recruitment of RNF34.

## Results

### RNF34 facilitates RGNNV replication though inhibiting IFN activation

To explore the role of RNF34 during NNV infection, we investigated the effect of RNF34 on NNV replication. As shown in Fig 1A and B, ectopic expression of RNF34 significantly increased CP expression, RNF34 knockdown resulted in a decreased transcription of NNV CP gene (Fig 1C and D). Furthermore, we found the transcription levels of IFNh and IFN-stimulated genes (ISGs), including ISG15, Viperin, and MX were significantly reduced by RNF34 (Fig 1E-H). Consistently, the result of luciferase reporter assays showed the IFNh promoter activity was significantly lower in RNF34 overexpressing cells than that in control group (Fig 1I), indicating that RNF34 might promote RGNNV replication by inhibiting IFN antiviral response.

**Fig. 1.**
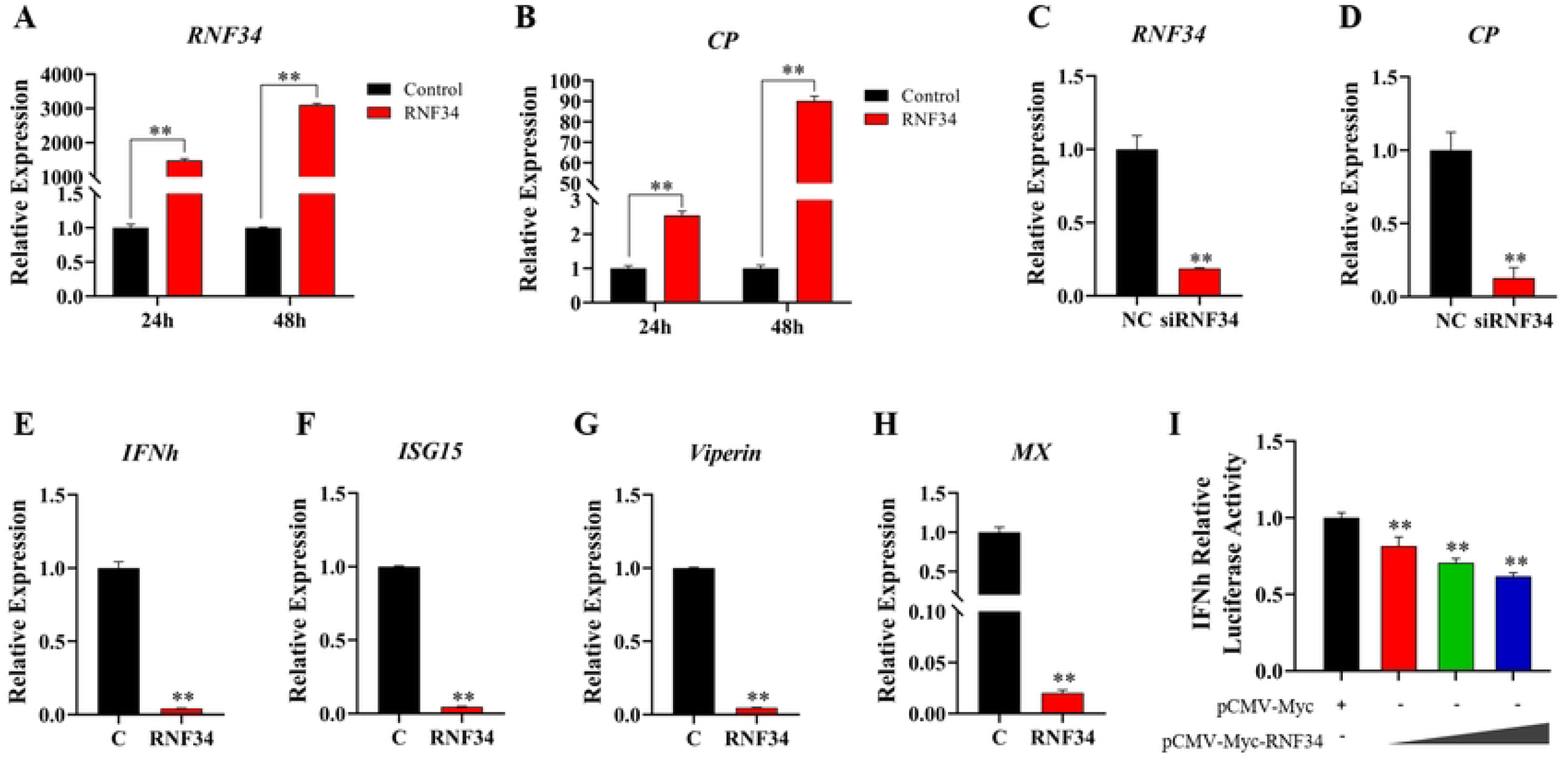
RNF34 promotes RGNNV infection and inhibits IFN responses. **(A-B)** LJB cells were transfected with *pCMV-Myc-RNF34* or *pCMV-Myc* plasmids (control), and infected with RGNNV for 24 h and 48 h, respectively. Then the cells were lysed for qRT-PCR to detect the expression of *RNF34* and *CP*. **(C-D)** qRT-PCR analysis of *RNF34* and *CP* mRNA expression in siRNF34 or NC (control) transfected LJB cells, following infection with RGNNV for 24 h. **(E-H)** qRT-PCR analysis of *IFNh, ISG15, Viperin* and *MX* expression in *pCMV-Myc-RNF34* transfected LJB cells, following infection with RGNNV for 24 h. **(I)** Luciferase activity of IFNh promoter in FHM cells transfected with an increasing amount of *pCMV-Myc-RNF34* plasmid (0, 100, 250, and 500 ng), together with reporter plasmids *pGL3-IFNh-pro-Luc* and renilla luciferase plasmid *pRL-TK*. Data is collected from three independent experiments and presented as mean ± S.D. * *p* < 0.05; ** *p* < 0.01.

### RNF34 interacts with TBK1 and IRF3 and inhibits TBK1 and IRF3-mediated IFN response

The RLR-induced IFN response is essential for fish innate immunity against NNV infection [19], thus, we firstly investigated whether RNF34 is a negative interactor of the key molecules in RLR signaling pathway. As shown in Fig 2A-D, RNF34 could be co-immunoprecipitated with both TBK1 and IRF3, but not with MAVS and TRAF3. Confocal microscopy analysis also showed both TBK1 and IRF3 were colocalized with RNF34 in the cytoplasm of HEK 293T cells (Fig 2E and F). Moreover, the interaction between RNF34 and TBK1 or IRF3 was also confirmed in another two model fish species, *Danio rerio* and *Oryzias melastigma* (Fig 2G and H). These data indicated that RNF34 universally interacted with TBK1 and IRF3.

**Fig. 2.**
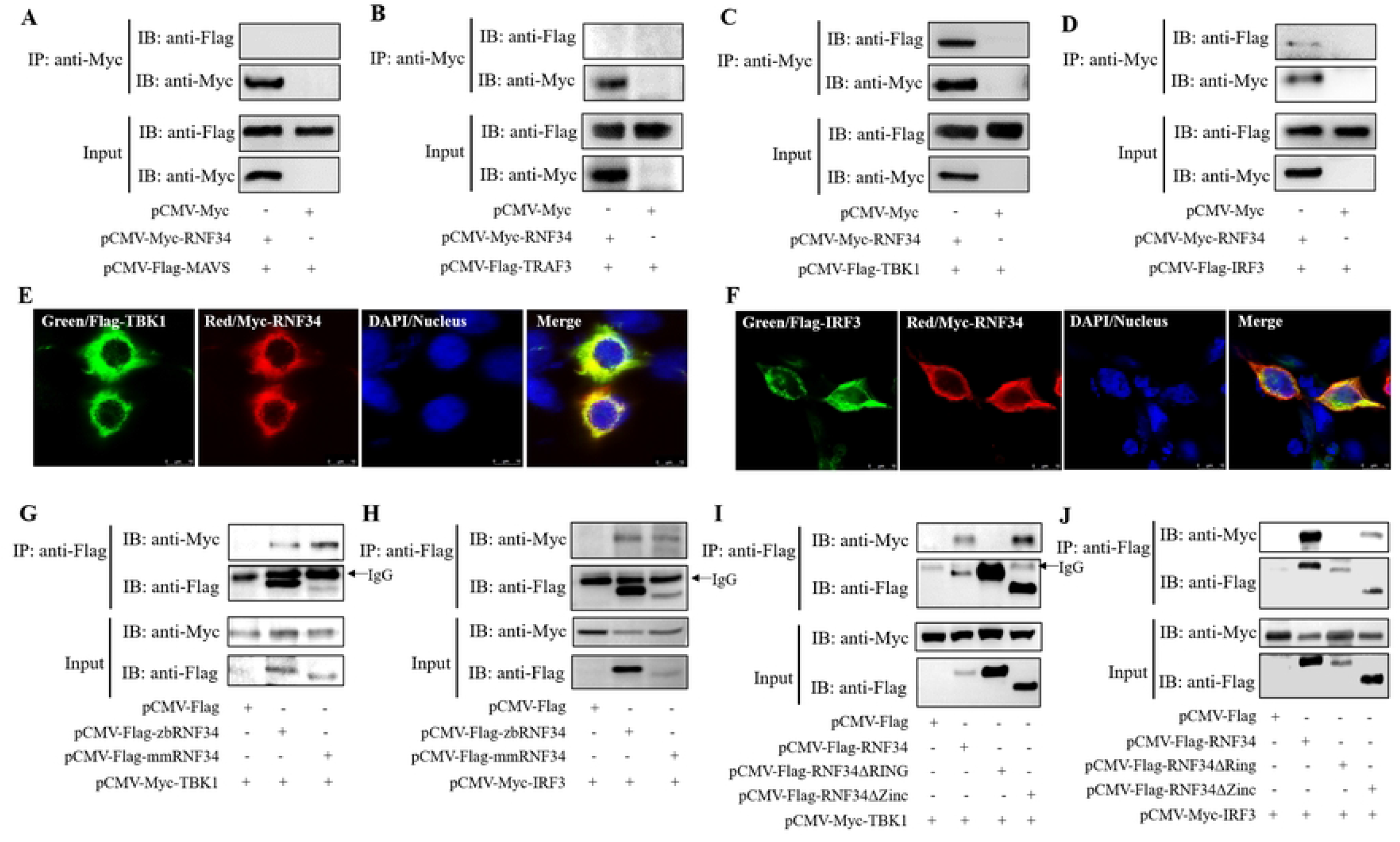
RNF34 interacts with TBK1 and IRF3 through RING domain. **(A-D)** HEK 293T cells were transfected with *pCMV-Myc-RNF34* and *pCMV-Flag-MAVS, pCMV-Flag-TRAF3, pCMV-Flag-TBK1* or *pCMV-Flag-IRF3* plasmids, respectively. At 24 h post transfection, the cell lysates were subjected to co-immunoprecipitation analysis with anti-Myc magnetic beads as indicated. **(E-F)** *pCMV-Myc-RNF34* and *pCMV-Flag-TBK1* or *pCMV-Flag-IRF3* plasmids were transfected into HEK 293T cells for immunofluorescence analysis by using anti-Myc (red) and anti-Flag (green) antibodies. Nuclei were stained with DAPI. **(G-H)** Plasmids of RNF34 from zebrafish (zbRNF34) and marine medaka (mmRNF34) were transfected into HEK 293T cells, together with *pCMV-Myc-TBK1* or *pCMV-Myc-IRF3* plasmids, respectively. At 24 h post transfection, the cell lysates were subjected to co-immunoprecipitation analysis with anti-Flag magnetic beads as indicated. **(I-J)** HEK 293T cells were transfected with Flag-Tagged RNF34 mutant plasmids (RNF34ΔRING and RNF34ΔZinc) as indicated for 24 h, the cell lysates were subjected to co-immunoprecipitation analysis with anti-Flag magnetic beads as above.

To map the key domain that mediated the binding of RNF34 to TBK1 and IRF3, a series of truncated RNF34 mutants were constructed. As shown in Fig 2I and J, Co-IP assays showed that the deletion of RING domain of RNF34 completely abrogated the interaction between RNF34 and TBK1 or IRF3, suggesting that RING domain is essential for their interaction.

To elucidate the effect of RNF34 on IFN response induced by TBK1 and IRF3, the luciferase reporter assay was conducted. As shown in Fig 3A and B, RNF34 had a dose-dependent inhibitory effect on the activation of the IFNh promoter mediated by TBK1 and IRF3. A domain mapping experiment further found that deletion of RING domain, but not the Zinc domain, lost the ability to suppress TBK1 and IRF3-induced IFNh promoter (Fig 3C and D). These data indicated that RNF34 negatively regulated RLR-induced host IFN response by targeting TBK1 and IRF3.

**Fig. 3.**
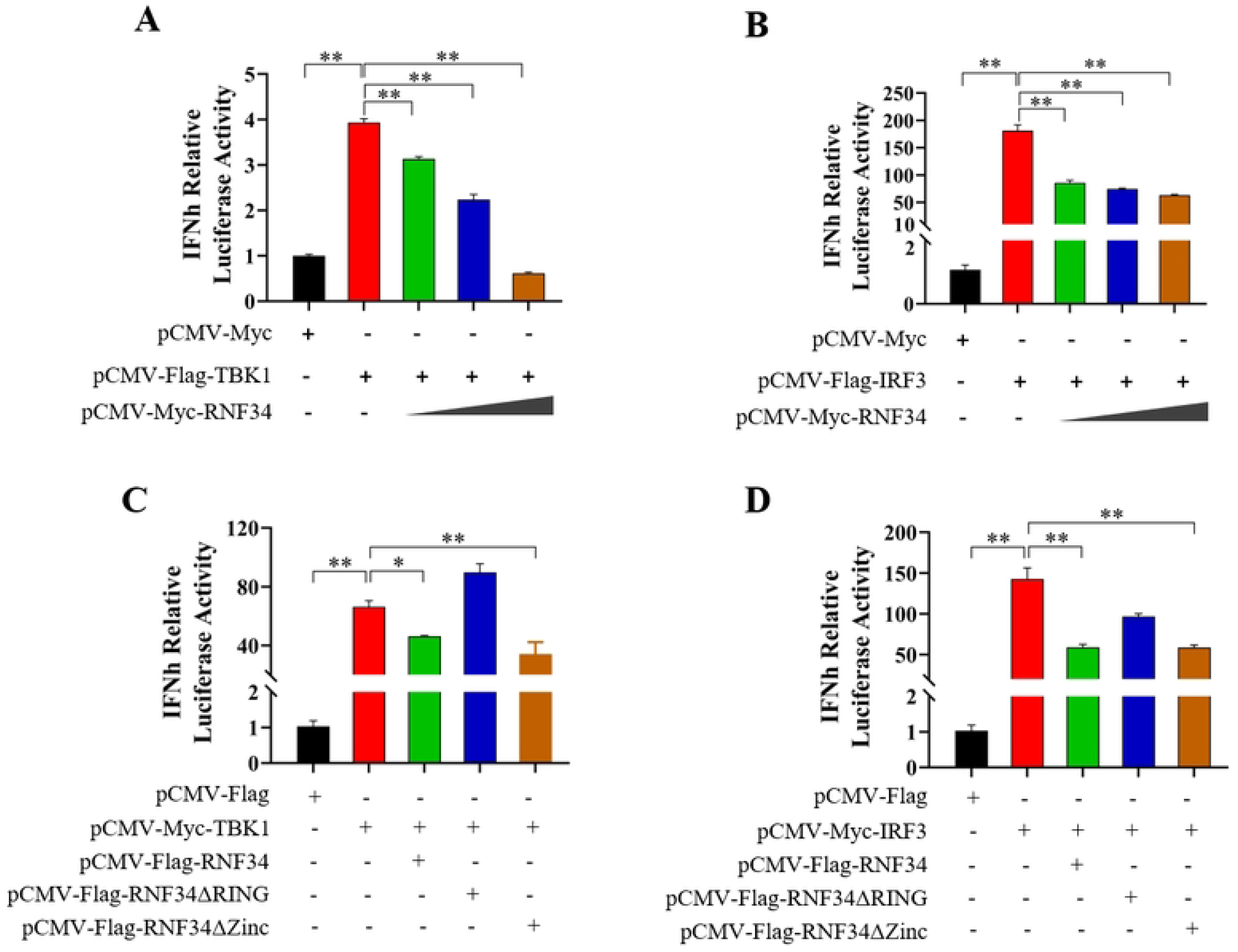
RNF34 inhibits TBK1 and IRF3-induced IFN responses. **(A-B)** FHM cells were co-transfected with an increasing amount of *pCMV-Myc-RNF34* (0, 100, 200, and 300 ng), *pGL3-IFNh-pro-Luc* and *pRL-TK*, together with *pCMV-Flag-TBK1* (A) or *pCMV-Flag-IRF3* (B) plasmids, respectively, for luciferase activity analysis. **(C-D)** RNF34 mutant plasmids were transfected into FHM cells for 24 h, along with *pCMV-Myc-TBK1* (C) or *pCMV-Myc-IRF3* (D) plasmids, respectively. Reporter assays were performed as above (n = 3). * *p* < 0.05; ** *p* < 0.01.

### RNF34 mediates K27- and K48-linked ubiquitination and degradation of TBK1 and IRF3

To elucidate the underlying mechanism of RNF34 on negative regulation of TBK1 and IRF3-mediated IFN response, the effect of RNF34 on TBK1 and IRF3 expression was investigated. RNF34 overexpression inhibited mRNA expression of both TBK1 and IRF3, whereas RNF34 knockdown led to the opposite effects (Fig 4A and B). Meanwhile, RNF34 decreased TBK1 or IRF3 protein levels in a dose-dependent manner in HEK 293T cells (Fig 4C and D). Consistently, RNF34 also reduced the protein expression levels of endogenous TBK1 and IRF3 in LJB cells without or with RGNNV challenge (Fig 4E and F). Furthermore, MG132 (a proteasome inhibitor) or NH_4_Cl (a lysosome inhibitor) was used to determine whether the proteasome or lysosome pathway was responsible for RNF34-induced TBK1 and IRF3 degradation. As shown in Fig 5A and B, MG132 restored RNF34 induced TBK1 and IRF3 degradation, but NH_4_Cl could not block the decrease of TBK1 and IRF3 caused by RNF34 (S1 Fig), indicating that TBK1 and IRF3 undergoes RNF34-mediated proteasomal degradation. Moreover, we found that RNF34 induced the K27- and K48-linked ubiquitination and degradation of TBK1 and IRF3 (Fig 5C and D). Consistently, luciferase assay results indicated that the inhibition effect of RNF34 on TBK1 and IRF3-induced IFNh reporter activation was enhanced by ubiquitin-K27 and ubiquitin-K48 (Fig 5E and F).

**Fig. 4.**
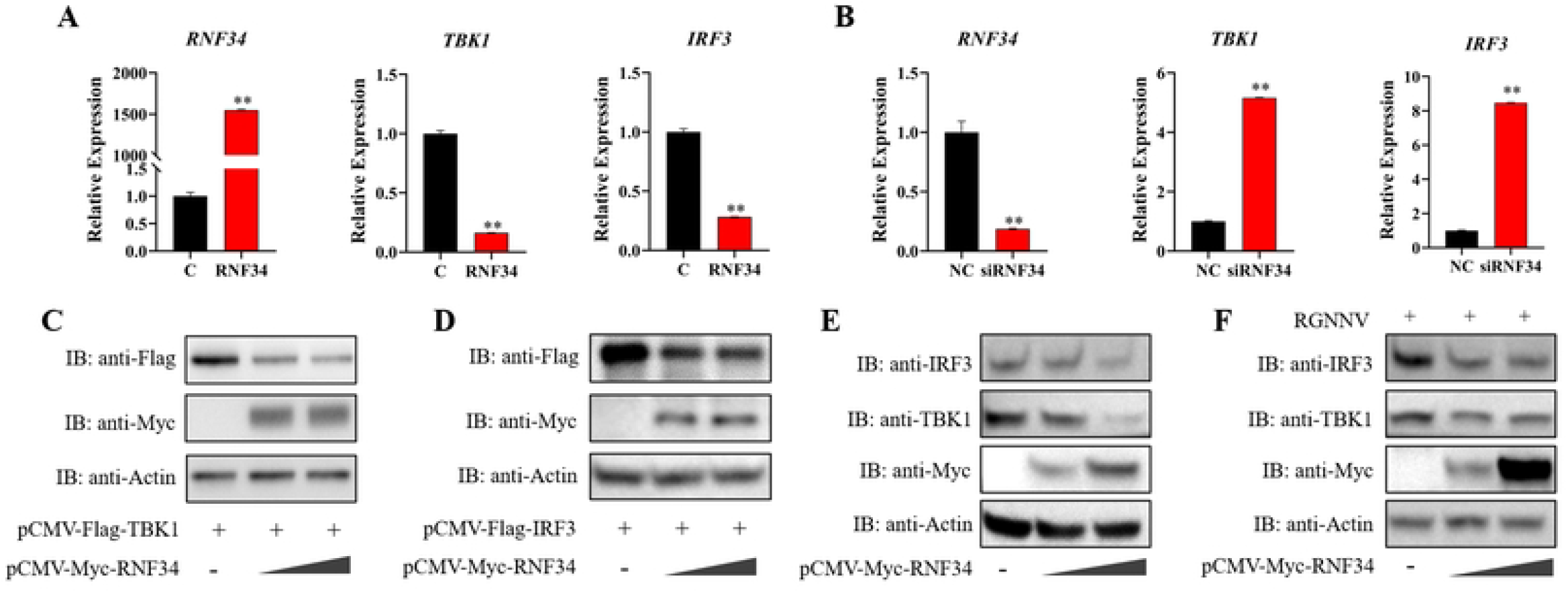
RNF34 enhances the degradation of TBK1 and IRF3. **(A-B)** qRT-PCR analysis of *RNF34, TBK1* and *IRF3* mRNA expression in LJB cells with *pCMV-Myc-RNF34* overexpressed (A) or RNF34-knock down (B), following infection with RGNNV for 24 h. **(C-D)** HEK 293T cells were transfected with the empty vector or *pCMV-Myc-RNF34* plasmid (0, 0.5, and 1 μg), together with *pCMV-Flag-TBK1* (C) or *pCMV-Flag-IRF3* (D) plasmids, respectively. At 24 h post transfection, the cells were lysed for immunoblot assays with indicated antibodies. **(E-F)** LJB cells transfected with *pCMV-Myc-RNF34* (0, 1.5, and 3 μg) without (E) or with RGNNV infection (F) were subjected to immunoblot assays using anti-TBK1 and anti-IRF3 antibodies.

**Fig. 5.**
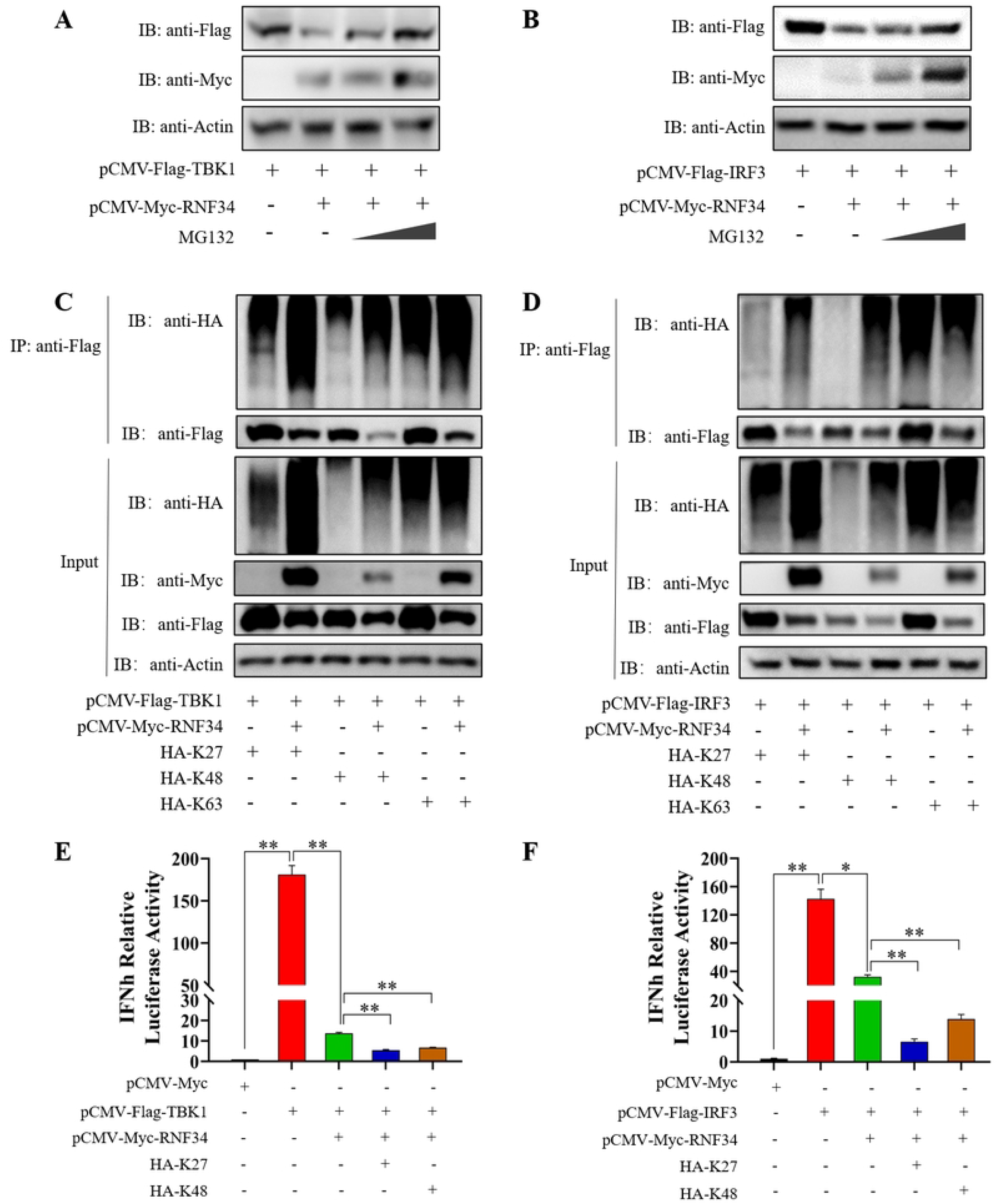
RNF34 promotes the K27 and K48-ubiquitination of TBK1 and IRF3. **(A-B)** HEK 293T cells were transfected with *pCMV-Myc-RNF34* plasmid, along with the *pCMV-Flag-TBK1* (A) or *pCMV-Flag-IRF3* (B) plasmids, and then stimulated with increasing amount of MG132 (10 and 20 μM) for 6 h. The cells were lysed for immunoblot assays with indicated antibodies. **(C-D)** HEK 293T cells were cotransfected with *pCMV-Myc-RNF34, HA-K27, HA-K48*, or *HA-K63*, along with the *pCMV-Flag-TBK1* (C) or *pCMV-Flag-IRF3* (D) plasmids for 24 h. Afterwards, the cells were lysed for co-immunoprecipitation analysis with anti-Flag antibodies as indicated. **(E-F)** Luciferase activity of IFNh promoter in FHM cells transfected with *pCMV-Myc-RNF34, HA-K27, HA-K48*, or *HA-K63*, along with the *pCMV-Flag-TBK1* (E) or *pCMV-Flag-IRF3* (F) plasmids for 24 h. Data is collected from three independent experiments and presented as mean ± SD. * *p* < 0.05; ** *p* < 0.01.

### RNF34 impairs TBK1-induced nuclear translocation of IRF3

Upon TBK1 activation, it would promote IRF3 translocating into the nucleus to activate the innate immunity and IFN production [20]. Thus, we further detected whether TBK1-induced IRF3 translocation was influenced by RNF34. As expected, when HEK 293T cells were transfected with TBK1 and IRF3, the cytoplasmic-localized IRF3 was observed in the nucleus; when the cells were further transfected with RNF34, IRF3 was predominantly found in cytoplasm, colocalized with the cytoplasmic RNF34 and TBK1 (Fig 6A). Consistent with the above observation, the Western blot analysis of cytoplasmic and nuclear fractions proved this finding, as TBK1 overexpression-induced the increasement of nuclear IRF3 protein was attenuated by RNF34 (Fig 6B).

**Fig. 6.**
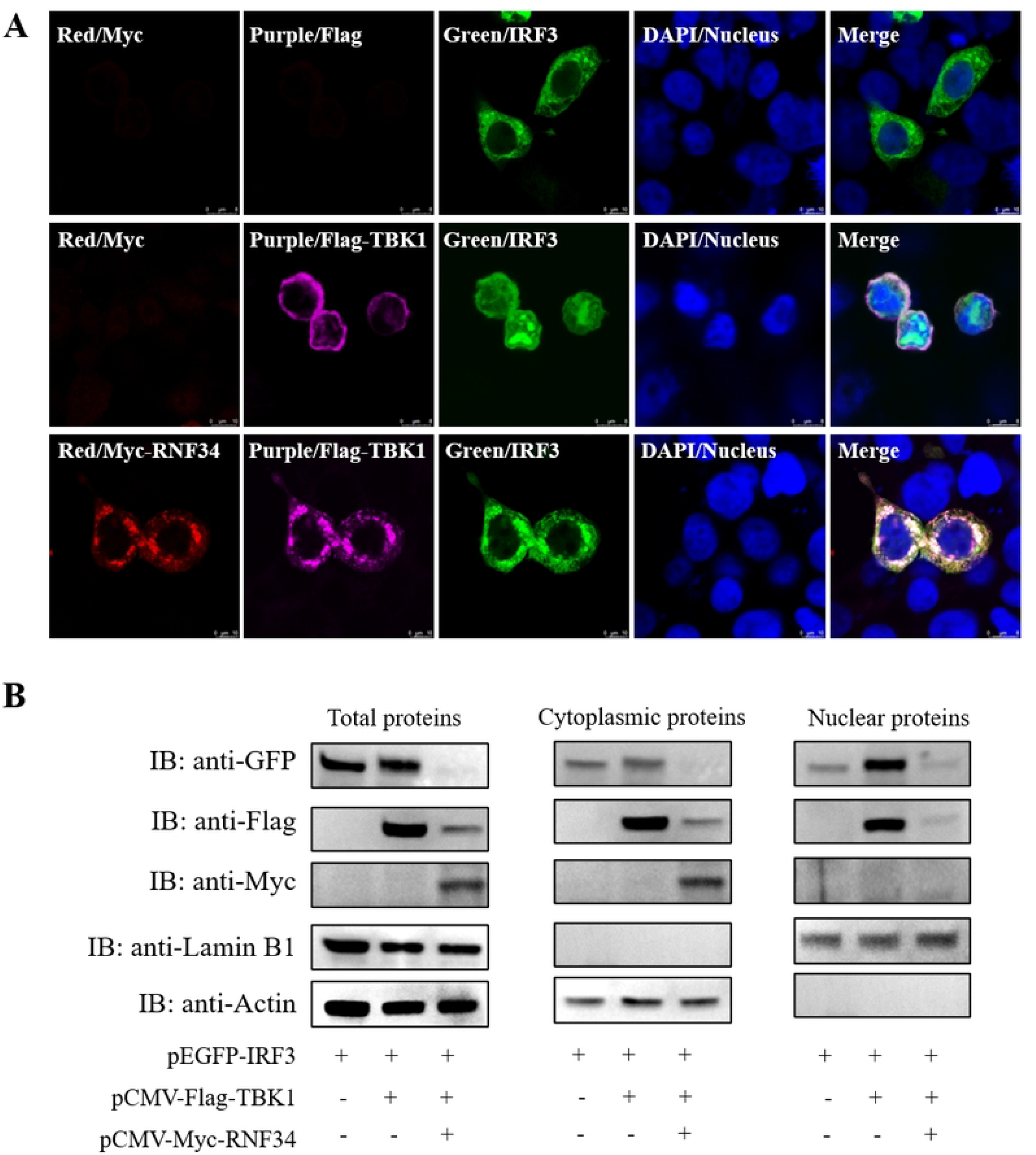
RNF34 diminishes TBK1-induced translocation of IRF3 from cytoplasm to nucleus. **(A)** *pCMV-Myc-RNF34* and *pCMV-Flag-TBK1* or *pEGFP-IRF3* plasmids were transfected into HEK 293T cells as indicated for immunofluorescence analysis by using anti-Myc (red) and anti-Flag (purple) antibodies. Nuclei were stained with DAPI. **(B)** HEK 293T cells were transfected with *pEGFP-IRF3* plasmid, along with *pCMV-Flag-TBK1* or *pCMV-Myc-RNF34* plasmids, then the cells were lysed for the cytoplasmic proteins, nuclear proteins and total proteins extraction and subjected to immunoblot assays using anti-GFP, anti-Flag, anti-Myc, anti-Lamin B1 and anti-Actin antibodies.

### CP recruits RNF34 to inhibit TBK1 and IRF3 induced IFN response

Our previous study has shown that CP can regulate protein ubiquitination to suppress RLR-mediated type-I IFN signaling [21]. Thus, we further examined the relationship between RNF34 and CP. Confocal microscopy and Co-IP assays revealed that RNF34 was associated with CP (Fig 7A-C). Pull-down assays showed a direct interaction between RNF34 and CP (Fig 7D). As shown in Fig 7E, the interaction relationship between RNF34 and CP was also confirmed in *Danio rerio* and *Oryzias melastigma*. In addition, we found that RNF34 was coprecipitated with CP wild-type, ARM domain deletion mutant, S domain deletion mutant, LR domain deletion mutant and P domain deletion mutant, but not with the arm domain deletion mutant (Fig 7F).

**Fig. 7.**
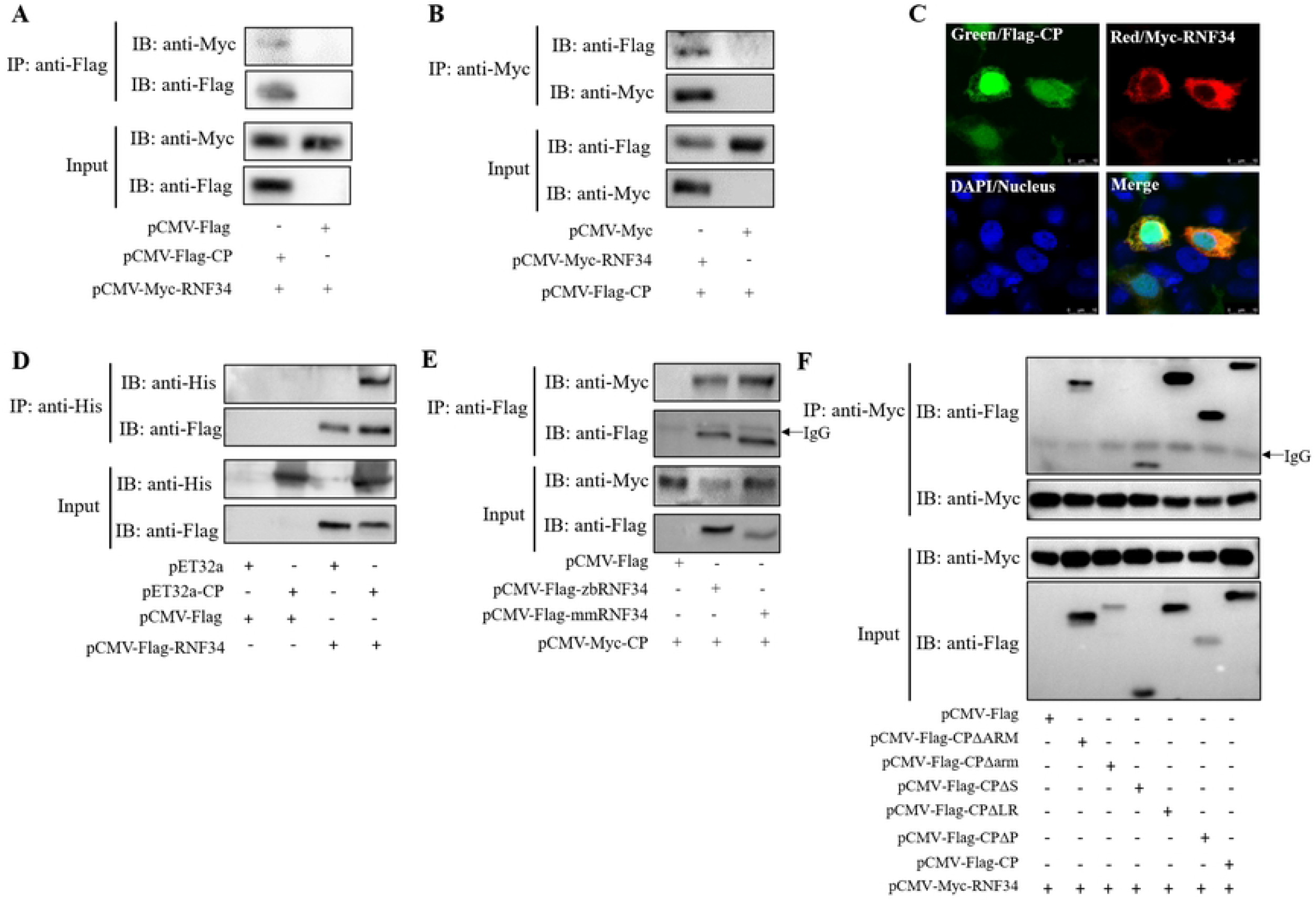
NNV capsid protein (CP) binds with RNF34. **(A-B)** HEK 293T cells were transfected with *pCMV-Myc-RNF34* and *pCMV-Flag-CP* plasmids as indicated for 24 h, the cell lysates were subjected to co-immunoprecipitation analysis with anti-Flag (A) or ant-Myc (B) magnetic beads as above. **(C)** His-CP or His proteins were purified to pull-down the protein lysates of HEK 293T cells transfected with *pCMV-Flag-RNF34* plasmids, the cell lysates were subjected to pull-down analysis with anti-His magnetic beads. **(D)** *pCMV-Myc-RNF34* and *pCMV-Flag-CP* plasmids were transfected into HEK 293T cells for immunofluorescence analysis by using anti-Myc (red) and anti-Flag (green) antibodies. Nuclei were stained with DAPI. **(E)** Plasmids of *pCMV-Flag-RNF34* from zebrafish and marine medaka were transfected into HEK 293T cells, together with *pCMV-Myc-CP* plasmids for 24 h, the cell lysates were subjected to co-immunoprecipitation analysis with anti-Flag magnetic beads as above. **(F)** HEK 293T cells were transfected with CP mutant plasmids (CPΔARM, CPΔarm, CPΔS, CPΔL, and CPΔP) as indicated for 24 h, the cell lysates were subjected to co-immunoprecipitation analysis with anti-Myc magnetic beads as above.

We further investigated the effect of CP on TBK1. CP induced the degradation of TBK1 in a dose dependent manner under endogenous and overexpressed conditions, and this effect could be recovered by MG132 treatment (Fig 8A-C), suggesting CP promoted the degradation of TBK1 through ubiquitination. Given the interaction between RNF34 and CP, we speculated that RNF34 might be involved in CP-induced TBK1 and IRF3 degradation. To test this hypothesis, we detected TBK1 and IRF3 expression in LJB cells cotransfected with CP plasmids and NC or siRNF34. The results showed that the inhibition of CP on TBK1 and IRF3 expression was decreased in the presence of siRNF34 in a dose-dependent manner, which would be further counteracted by RNF34 ectopic expression (Fig 8D). Consistently, luciferase reporter assays showed that siRNF34 attenuated CP-reduced IFNh promoter activity (Fig 8E). Taken together, these results demonstrated that CP utilized RNF34 to reduce IFN production via promoting TBK1 and IRF3 ubiquitination and degradation.

**Fig. 8.**
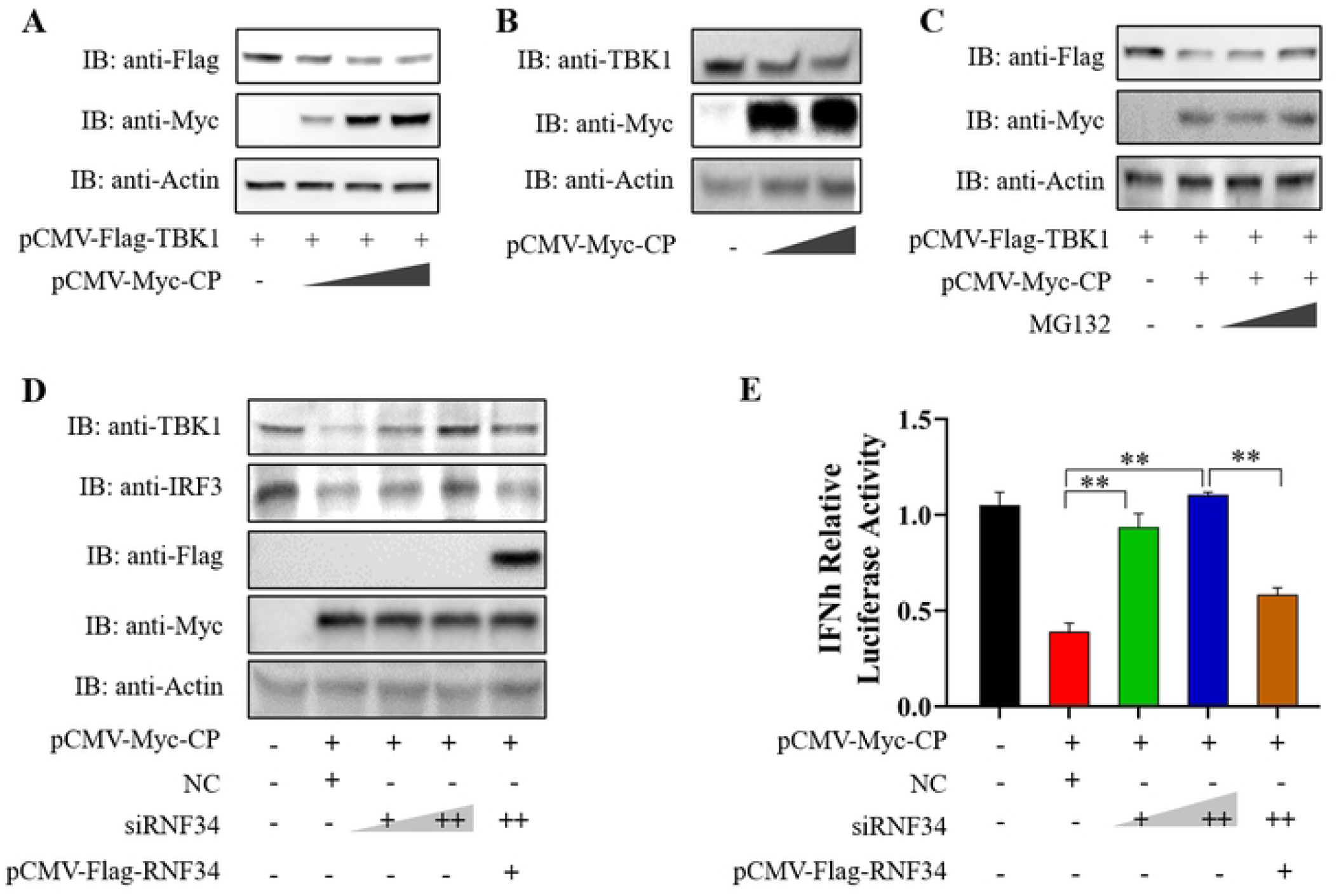
CP induces TBK1 and IRF3 degradation and IFN suppression depended on RNF34. **(A)** HEK 293T cells were transfected with the *pCMV-Flag-TBK1* and *pCMV-Myc-CP* plasmids (0, 0.5, and 1 μg) for 24 h, the cells were then lysed for immunoblot assays with indicated antibodies. **(B)** LJB cells transfected with *pCMV-Myc-CP* plasmid (0, 0.5, and 1 μg) for 24 h, and then were subjected to immunoblot assays using anti-TBK1 antibodies. **(C)** HEK 293T cells were transfected with *pCMV-Flag-TBK1* and *pCMV-Myc-CP* plasmids, followed by stimulation with increasing amount of MG132 (10 and 20 μM) for 6 h, the cells were lysed for immunoblot assays with indicated antibodies. **(D)** HEK 293T cells were transfected with *pCMV-Myc-CP* plasmids, along with the NC or siRNF34 (50 and 100 nm), or 100 nm siRNF34 plus *pCMV-Flag-RNF34* plasmids. At 24 h post transfection, the cell lysates were subjected to immunoblot assays with indicated antibodies. **(E)** Luciferase activity of IFNh promoter in FHM cells transfected with *pCMV-Myc-CP* plasmids, along with the NC or siRNF34 (50 and 100 nm), or 100 nm siRNF34 plus *pCMV-Flag-RNF34* for 24 h. Data is collected from three independent experiments and presented as mean ± SD. * *p* < 0.05; ** *p* < 0.01.

## Discussion

To survive in hosts, viruses have evolved various strategies to evade host antiviral innate immunity for their replication. As a pivotal sensor of RNA viruses and activator of IFN production, RLRs-mediated signaling is tightly regulated by host and viral factors [22]. Here, we identified that NNV evaded RLRs-mediated IFN response via the host E3 ubiquitin ligases RNF34. Mechanistically, NNV CP blocked the RLRs signaling pathway by binding with RNF34 for ubiquitination degradation of TBK1 and IRF3 (Fig 9). RNF34, a caspase 8/10-associated ubiquitin ligase, was firstly identified as a RING-type E3 ubiquitin ligase in human [23]. Emerging evidence has shown that RNF34 is involved in many biological processes, such as the development of multiple neurological disease, brown fat cell metabolism and immune response [24]. For example, RNF34 inhibited activation of NF-κB through direct interaction and ubiquitination of NOD1 [25]. A recent study showed that RNF34 negatively regulated RLRs-mediated antiviral immunity responses by promoting autophagic degradation of MAVS [26]. Hence, our finding demonstrates a novel distinct mechanism of RNF34 functioning as an IFN antagonist via targeting TBK1 and IRF3 for ubiquitination and degradation, which highlights the importance of RNF34 in regulation of RLRs-mediated signaling pathway.

**Fig. 9.**
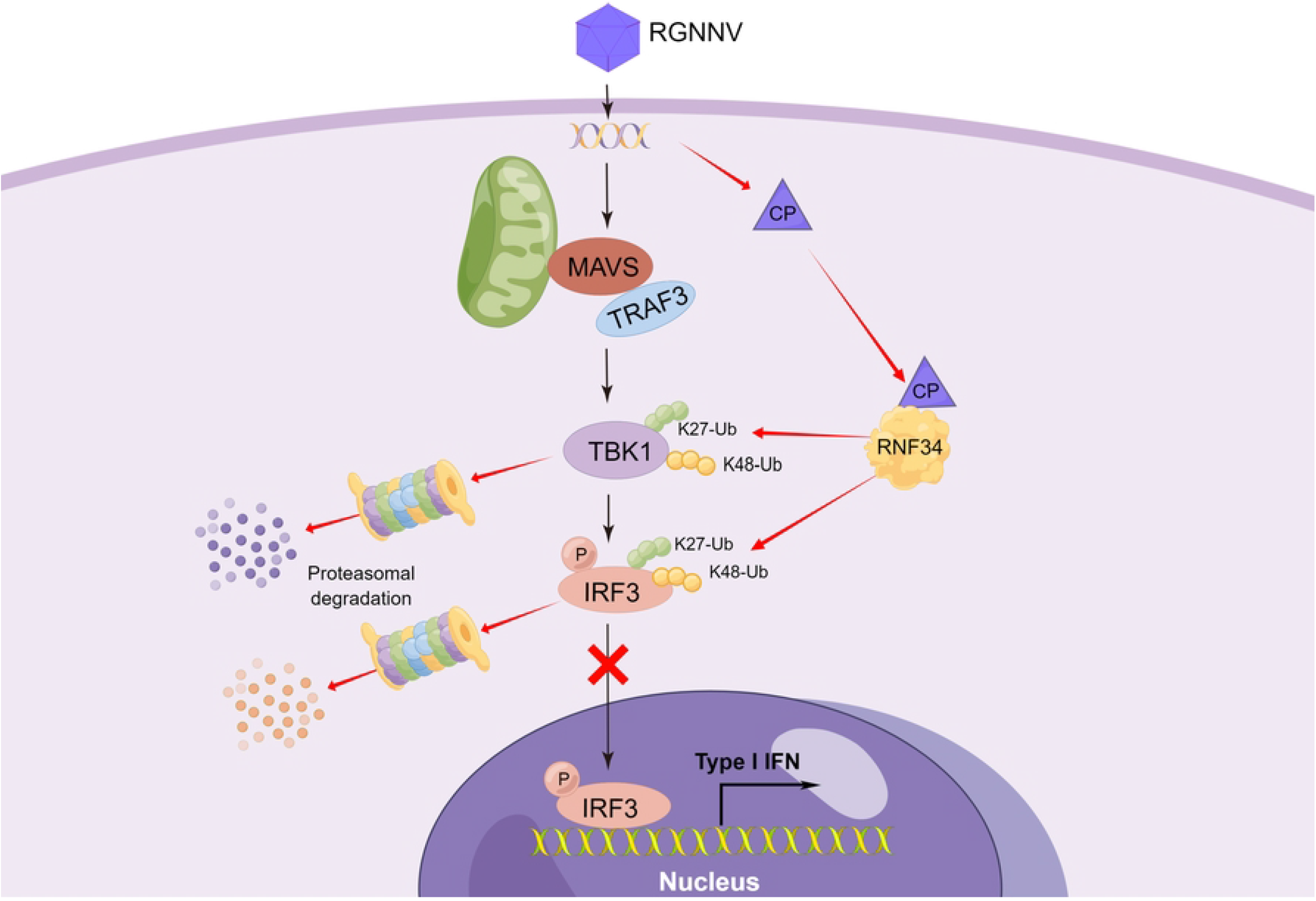
A proposed working model of NNV evades antiviral innate immunity by inhibiting RIG-I-like receptor-mediated signaling via RNF34. CP interacted with RNF34 and utilized RNF34 to promote K27- and K48-linked ubiquitination and degradation of TBK1 and IRF3, thus impeding the translocation of IRF3 into the nucleus, finally suppressing the production of IFN. Schematic figure was drawn by Figdraw.

Ubiquitination is an essential posttranslational modification that plays crucial roles in the control of antiviral immunity upon virus infection. As crucial components of the ubiquitination system, an increasing number of RING-domain E3 ligases have emerged as key regulators of immune responses. For instance, RNF122 promoted RIG-I degradation via K48-linked ubiquitination to inhibit host immunity against virus infection [12]. RNF19a catalyzed K48-linked ubiquitination to degrade RIG-I, and finally attenuated RIG-I-mediated immune responses [27]. Here, we found that RNF34 functions as a negative regulator of RLRs-mediated signaling pathway to facilitate NNV replication via targeting TBK1 and IRF3. Importantly, the interaction between RNF34 and TBK1 or IRF3 was conserved in different fishes, indicating the conservation of RNF34’ function. The RING domain is the critical functional domain for RING finger family E3 ubiquitin ligases. Truncation mutation within RING domain abolished the ability of Mex3A to induce the ubiquitination and degradation of RIG-I [28]. TRIM40 mutant lacking RING domain greatly impaired its inhibition on IFN-b promoter activity [5]. In this study, we found that RNF34 interacted with TBK1 and IRF3 via its RING domain, and the RING domain is indispensable for the ability of RNF34 to restrain IFN signaling, indicating that the E3 ligase activity of RNF34 is essential for its regulatory function on TBK1 and IRF3.

TBK1 and IRF3 are important adaptors of the RLRs-mediated signaling pathway, therefore, both are tightly regulated by a variety of mechanisms such as ubiquitination, phosphorylation, prevention of active TBK1 complexes formation, and blocking of the IRF3 translocation from cytoplasm into the nucleus [27]. Recently, emerging evidence has shown the regulation of TBK1-IRF3 activity via the ubiquitin system. For example, multiple E3 ubiquitin ligases, such as Triad3A [29], TRIM27 [30] and DXT4 [4] target TBK1 for K48-linked polyubiquitination degradation, thus negatively regulating type I IFN production. TRIM26 [31], RAUL [32], and RBCC protein interacting with PKC [33], conjugate K48-linked polyubiquitin chains on IRF3, resulting in proteasomal degradation of IRF3 and inhibition of host antiviral innate immune response. Here, RNF34 was found to mediated K27 and K48-linked polyubiquitination of both TBK1 and IRF3, thus inhibit IFN signaling. In addition, we also found that RNF34 not only led to the degradation of IRF3, but the inhibition of TBK1-induced IRF3 nuclear translocation. Considering the importance of IRF3 nuclear translocation for the activation of the IFN-I promoter, we speculated that RNF34 interfered with TBK1-induced IRF3 nuclear translocation by degrading TBK1, subsequently dampened type I IFN response. To the best of our known, only MAVS had been identified as a target of RNF34 negatively regulating RLRs-mediated antiviral immunity. Hence, our study provides novel target molecules of RNF34 in RLRs-mediated signaling pathway.

Given the important role of ubiquitination in the innate immune response, many viruses have evolved elaborate mechanisms to directly or indirectly hijack the host ubiquitin system to favor self-replication. For instance, the V protein of Newcastle disease virus inhibited IFN signaling by promoting ubiquitination-dependent degradation of MAVS via RNF5 [34]. African swine fever virus pI215L protein recruited RNF138 to reduce K63-linked ubiquitination of TBK1, resulting in the inhibition of IFN production [35]. The NNV CP is responsible for host innate immune evasion, however its exact evasion mechanisms are not well characterized. We and others have previously shown that the ubiquitin proteasome system played an important role during NNV infection [17, 36]. Recently, we found NNV CP targeted sea perch TRAF3 and IRF3 to promote their ubiquitination degradation, leading to the inhibition of IFN response. Of note, RNF114 was exploited by CP through its P domain to potentiates K27- and K48-linked ubiquitination of TRAF3 [21, 36]. However, the E3 ubiquitin ligase participated in CP-induced IRF3 ubiquitination degradation was still unknown. Here, we found that CP upregulated the expression of RNF34 and directly interacted with RNF34, indicating that RNF34 might be associated with CP induced IRF3 degradation. Importantly, their interaction is conservative in different fishes. CP consists of four domains and a linker region (L) [37]: the N-terminal ARM (ARM), which contains a nucleolus localization signal (aa 23 to 31) associating with cell cycle arrest; N-terminal arm (arm), a conserved region that recruits the RNA during encapsidation; the shell domain (S), which is responsible for virus assembly, and the protrusion domain (P) that is involved in interacting with the host cell surface and the trimerization of the protein [16]. Interestingly, the diverse CP domains had been found responsible for different conjugated protein localization, such as NNV receptor HSC70 bind on ARM domain; HSP90ab1 target on L domain [38]; the CP S domain is contributed to interactions with both of IRF3 and TRAF3 [21]. Here, different with them, domain mapping showed RNF34 bind to the arm domain of CP, implying a novel molecular function of CP arm domain.

Furthermore, our data demonstrated that NNV CP could induce TBK1 degradation, which can be recovered by MG132 treatment and deficiency of RNF34, suggesting that RNF34 was responsible for CP-mediated TBK1 degradation. Meanwhile, deficiency of RNF34 also impaired CP-mediated IRF3 degradation. All these results demonstrated that CP recruited RNF34 to mediate TBK1 and IRF3 ubiquitination and degradation. Previously, we had reported that CP hijacked RNF114 to catalyze the K27- and K48-linked ubiquitination of TRAF3 for proteasomal degradation, decreasing TRAF3-mediated IFN signaling [21]. The findings identified a novel strategy adopted by NNV to evade host antiviral innate immunity via hijacking the RNF proteins-mediated ubiquitination process and suggested that CP could utilize multiple E3 ubiquitin ligases to suppress the RLRs-mediated antiviral signaling. Considering the interactions of CP and RNF34, RNF34 and TBK1 or IRF3 are conserved in different fishes, we speculate that it is a general immune evasion strategy exploited by CP to target against the IFN response via RNF34.

Overall, we have identified RNF34 as a suppressor of host innate immune response to promote NNV infection. Furthermore, NNV CP hijacks RNF34 to induce the ubiquitination and degradation of TBK1 and IRF3 for inhibition of IFN signaling. These findings reveal a new mechanism used by CP to counteract the IFN responses for supporting viral proliferation and provide a new understanding of the immune evasion strategies used by NNV.

## Materials and Methods

### Cell culture and reagents

LJB cells derived from sea perch (*Lateolabrax japonicus*) brain were cultured in DMEM medium (Gibco) with 15% FBS at 28°C [39]. Fathead minnow (FHM) cells were cultured in M199 medium (Gibco) with 10% FBS at 28°C. Human embryonic kidney 293T (HEK 293T) cells were maintained in DMEM/F12 with 10% FBS at 37°C in 5% CO_2_ hatchery.

Antibodies to Flag tag (M20008L), Myc tag (M20002L), His tag (M20001L), HA tag (M20003L) and actin (P30002L) were obtained from Abmart (Guangzhou, China). Antibodies to TBK1 (bs-7497R) and IRF3 (bs-1185R) were obtained from Bioss (Beijing, China). Anti-Lamin B1 antibodies (CPA1693) were purchased from Cohesion Biosciences (Shanghai, China). Donkey anti-mouse or goat anti-rabbit IgG (H + L) highly cross-adsorbed secondary antibodies, Alexa Fluor™ 555 and Alexa Fluor™ 488 were obtained from Invitrogen (Carlsbad, CA, USA). Magnetic beads of anti-Flag (HY-K0207), anti-c-Myc (HY-K0206) and anti-His (HY-K0209) were purchased from MedChemExpress (Monmouth Junction, NJ, USA). MG132 (M7449), NH_4_Cl (A9434), DAPI (D9542), phenylmethylsulfonyl fluoride (P7626), and isopropyl-1-thio-β-D-galactopyranoside (IPTG) (I6758) were procured from Sigma-Aldrich (St. Louis, MO).

### Plasmids construction

The full-length sequences of sea perch *RNF34* (GenBank accession number: OP784387) was amplified by PCR using primers (S1 Table) and was subsequently cloned into *pCMV-Flag/Myc* vector (Clontech). RNF34 deletion mutants (RNF34ΔRING and RNF34ΔZinc) were constructed by PCR and subcloned into *pCMV-Flag* vector. RNF34 from zebrafish (*Danio rerio*) and marine medaka (*Oryzias melastigma*) were cloned into *pCMV-Flag* vector, respectively. *Flag-MAVS, Flag-TRAF3, Flag-IRF3, Flag-TBK1, Myc-IRF3, Myc-TBK1, pGL3-IFNh-pro-Luc, pRL-TK, HA-K27, HA-K48, HA-K63, Flag-CP, pET32a(+)-CP* and truncated mutants of CP with Flag tags were obtained as described previously [21].

### Cell transfection and NNV infection

LJB cells in six-well plates (1 × 10^6^ cells/well) were transfected with different plasmids using Lipofectamine 8000 (Beyotime, Shanghai, China) following the manufacturer’s instructions. Post 24 h transfection, the cells were infected with red-spotted grouper NNV (RGNNV) at a multiplicity of infection (MOI) of 1 for the indicated hours and examined by quantitative reverse transcription-PCR (qRT-PCR) or Western blot.

### Cells stimulation

For stimulation, the cells in six-well plates were treated with proteasomal inhibitor MG132 (20 or 50 mM), lysosomal inhibitor NH_4_Cl (20 or 50 mM) or DMSO for 6 h, respectively. Post 24 h transfection, the cells were subjected to Western blot analysis.

### RNA interference

Small interfering RNAs (siRNAs) targeting RNF34 (siRNF34) were synthesized by the Ribobio company (Guangzhou, China), including siRNA-1: 5’-CACCGATACCTGCAGGGA-3’; siRNA-2: 5’-GAGGAAGAGGAGGACCC-3’; siRNA-3: 5’-AAGAACAGGAAATCATT-3’; and control siRNA (NC): 5’-UUCUCCGAACGUGUCACGUTT-3’. Cells were transfected with the indicated siRNA as described previously [40].

### qRT-PCR

Total RNA of cultured cells was extracted with TRIzol reagent (Invitrogen, CA, USA) according to the manufacturer’s instructions and was further reverse-transcribed into cDNA through the PrimeScript™ RT Reagent Kit with gDNA Eraser (TaKaRa). A LightCycler 480 II (Roche Applied Science, Germany) and SYBR RT-PCR kit (Roche) were used for qRT-PCR analysis by using gene-specific primers (S1 Table). mRNA relative expressions were evaluated from triplicate experiments and normalized to sea perch *β-actin*. The relative fold induction of genes was calculated using the 2^−ΔΔCt^ method and presented as mean ± S.D.

### Western blot and Co-immunoprecipitation (Co-IP)

The cells were lysed with lysates buffer (Beyotime), and boiled 10 min with 1% SDS for SDS-PAGE separation. The proteins were transferred onto PVDF membranes (Millipore, USA), then blocked with 5 % non-fat dried milk for 1 h at room temperature (RT), followed by primary antibodies incubation at 4 °C overnight, including anti-Flag (1:4000), anti-Myc (1:4000), anti-HA (1:4000), anti-TBK1 (1:1000), anti-IRF3 (1:1000), anti-actin (1:2000) or anti-Lamin B1 (1:1000) antibodies. The membranes were further probed with donkey anti-mouse or goat anti-rabbit IgG (H + L) highly cross-adsorbed secondary antibodies (1:1000) for 1 h at RT, and analyzed using ECL immunoblotting detection reagents (Millipore, USA) on a chemiluminescence instrument (Sage Creation, China).

For Co-IP assay, cell extracts were incubated with anti-Flag/Myc magnetic beads at 4 °C overnight. The beads were washed five times with lysis buffer and eluted with 1% SDS buffer for boiling and Western blot.

### Pull-down

Pull-down assays were performed as described previously [38]. His-fused CP proteins were extracted from *pET-32a (+)-CP* plasmids-transformed *E. coli BL21*(DE3) cells with 0.5 mM IPTG stimulation. His-Tag magnetic beads were firstly mixed with the His-fused CP proteins for 4 h at RT. Then the beads were incubated with protein lysates from HEK 293T cells post *pCMV-Myc-RNF34* transfection at 4 °C overnight, and finally analyzed by Western blot.

### Immunofluorescence assays

HEK 293T cells plated on coverslips in 24-well plates were transfected with different plasmids. After 24 h transfection, cells were washed twice with PBS and fixed with 4% paraformaldehyde for 1 h. After permeabilization with 0.15% Triton X-100 for 10 min and blocking with 5% skim milk for 1 h, the cells were incubated with primary antibodies at 4 °C overnight, including anti-Flag (1:500) and anti-Myc (1:500) antibodies, and followed by incubation with Alexa Fluor™ 555 or 488 conjugated secondary antibodies against mouse IgG (1:1000). The coverslips were washed with PBS and observed under a SP8 Leica laser confocal microscopy imaging system (Leica, Germany).

### Luciferase reporter assays

FHM cells in 24-well plates were transfected with the sea perch IFNh promoter luciferase reporter plasmid (*pGL3-IFNh-pro-Luc*) and indicated plasmids. A renilla luciferase plasmid (*pRL-TK*) was co-transfected as an internal control. After 24 h transfection, the cells were lysed, the luciferase activity in cells was analyzed using a GloMax 20/20 luminometer with the Dual-Luciferase Reporter Assay system (Promega).

### Statistical analysis

Data are collected from three independent experiments, analyzed through SPSS version 20.0, and presented as the means ± S.D. Student’s t-test or one-way ANOVA was used for the statistical comparisons between two-group or multiple group comparisons, respectively. *p* < 0.05 was considered with statistically significant difference; *p* < 0.01 was considered with highly significant difference.

## Supplementary information

**S1 Fig. RNF34-mediated degradation of TBK1 and IRF3 is not affected by NH**_**4**_**Cl**. HEK 293T cells were transfected with *pCMV-Myc-RNF34* plasmids, along with the *pCMV-Flag-TBK1* **(A)** or *pCMV-Flag-IRF3* **(B)** plasmids, and then stimulated with increasing amount of NH_4_Cl (10 and 20 μM) for 6 h, the cells were lysed for immunoblot assays with indicated antibodies.

**S1 Table. The sequence and PCR efficiency of primers used in this study.**

